# Next-Generation saRNA Platforms: Systematic Screening and Engineering Enhances Superior Protein Expression and Organ-Specific Targeting for RNA Therapeutics

**DOI:** 10.1101/2025.03.30.644708

**Authors:** Zhen Sun, Yuxiao Liu, Haoyi Zhang, Ting Ge, Yuting Pan, Yang Liu, Miaomiao Wu, Tao Shan, Guoqiang Zhu, Qi Wu, Kangming Chen

## Abstract

Self-amplifying RNA (saRNA), derived from alphaviruses, encodes nonstructural proteins (NSP1-4) that form a replicase complex, enabling prolonged and amplified protein expression at lower doses than conventional mRNA. However, its high immunogenicity and limited replication efficiency require optimization for improved protein expression. Here, we systematically optimized a widely used Venezuelan Equine Encephalitis Virus (VEEV)-based saRNA by refining its capping structure, nucleotide modifications, polyA tail length, and regulatory elements, achieving over 10-fold enhancement in protein expression compared to the wild-type. Additionally, we screened 28 alphavirus-derived saRNA constructs in vitro, selecting the top 3 candidates Everglades virus (EVEV), Mosso das Pedras virus (MDPV), and Rio Negro virus (RNV) for in vivo evaluation using SM102 lipid nanoparticle (LNP) formulations and luciferase reporters. These optimized saRNAs demonstrated sustained expression and distinct extrahepatic tissue-specific biodistribution, notably high spleen targeting compared to conventional mRNA. Such organ-specific expression profiles present novel strategies for targeted RNA therapeutics, and further integration with advanced delivery technologies could enhance the precision, efficacy, and therapeutic potential of mRNA-based treatments.

TOC figure

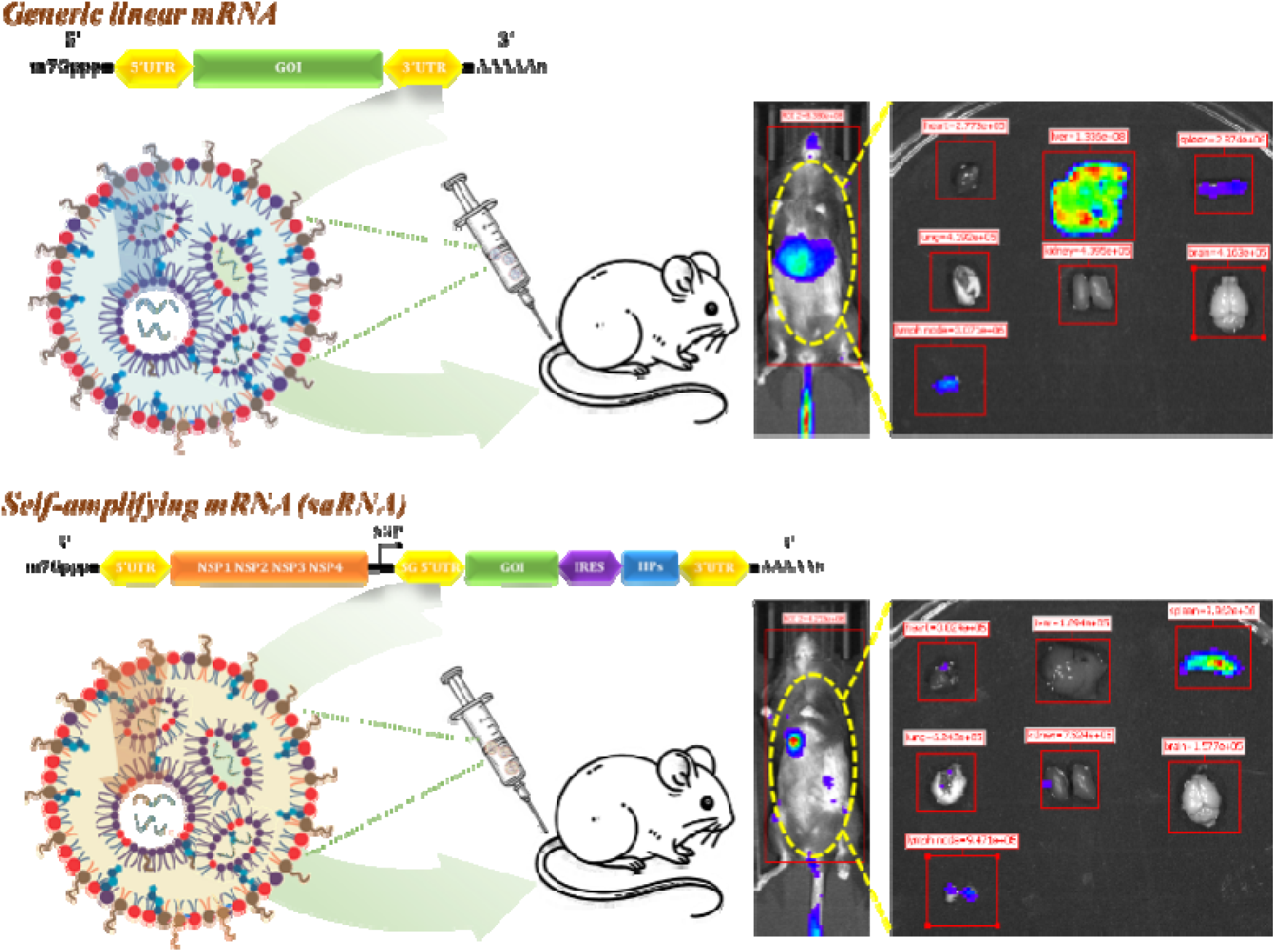

## Introduction

In recent years, rapid advancements in mRNA technology have significantly reshaped modern pharmaceutical science, providing simplified manufacturing processes, efficient purification, and improved scalability compared to traditional therapeutic approaches^1–3^. However, despite these advantages, instability, rapid degradation, and short half-life of conventional mRNA remain critical challenges, driving the need for optimized RNA design and delivery strategies^4–7^. Among these novel RNA approaches, self-amplifying RNA (hereinafter referred to as saRNA), derived from positive-sense alphaviruses, represents a promising platform. The saRNA vectors, including the wildly used Venezuelan Equine Encephalitis Virus (VEEV), encode viral nonstructural proteins (NSP1-4) that form an RNA-dependent RNA polymerase complex, enabling continuous amplification and prolonged protein expression from relatively low doses compared to conventional linear mRNA^8–12^. The recent approval of the saRNA-based COVID-19 vaccine ARCT-154 in Japan (2023) and the EU (2025) highlights its clinical relevance and potential^13,14^. Despite these advancements,, saRNA replication inherently generates double-stranded RNA (dsRNA) intermediates, activating host innate immune pathways such as RIG-I/MDA5 and PKR, thereby suppressing translation efficiency and reducing target protein expression^15–17^.

To optimize saRNA efficacy while minimizing unwanted immune activation, researchers have investigated various approaches, including sequence optimization, co-expression of immune-modulating proteins, and selection of replicon variants with reduced immunogenicity^15,18–20^. For example, Sahin et al. successfully employed co-delivery of immune evasion proteins (E3/K3/B18) to inhibit innate immune responses, significantly improving saRNA expression in vivo^21^. Likewise, Li et al. utilized an in vitro evolution method to select mutant Venezuelan Equine Encephalitis Virus (VEEV) replicons exhibiting enhanced protein expression and lower immune stimulation^22^.

Beyond these optimization methods, alphaviruses themselves represent a valuable source for enhancing saRNA performance due to their inherent diversity and tissue-specific tropisms. Alphaviruses comprise over 30 distinct subtypes, including extensively studied members such as Semliki Forest Virus (SFV), Sindbis Virus (SINV), Eastern Equine Encephalitis Virus (EEEV), and VEEV^23^. These viruses exhibit natural tropisms for specific tissues—VEEV preferentially infects neurons and lymphoid tissues, SINV targets muscle and skin cells, and Chikungunya virus (CHIKV) has affinity for fibroblasts and joint tissues^24–26^. Based on these inherent viral tropisms, it is hypothesized that saRNAs derived from distinct alphaviruses may similarly possess unique organ-specific expression patterns. Currently, targeted delivery of mRNA therapeutics primarily relies on modifying lipid nanoparticle (LNP) formulations, such as ionizable lipids screening and different ligand (e.g. peptide, antibody) conjugation^27–29^. These representative targeting strategies emphasizes the design and optimization of delivery vehicles. Alternatively, leveraging the intrinsic tropisms of alphavirus-derived saRNAs could offer a cargo-based complementary strategy for targeted RNA delivery.

Despite the widespread use of VEEV saRNA, there remains a need for a more effective saRNA vector that offers enhanced protein expression, low immunogenicity and distinct tissue-specific expression patterns. In this study, we aimed to optimize the design of saRNA vector and systematically screen alphavirus-derived saRNA constructs to identify sequences exhibiting enhanced expression efficiency and desirable tissue-specific expression profiles. Initially, the VEEV-based saRNA construct was optimized, yielding a variant with significantly improved protein expression (>10-fold over wild-type). Subsequently, a library containing 28 distinct alphavirus-derived saRNA vectors was constructed and screened in vitro. The top-performing saRNAs were further evaluated in vivo, using a luciferase reporter system to characterize their biodistribution and tissue tropisms. For the first time, our results showed that selected saRNAs exhibited sustained protein expression with unique organ-specific distribution, providing a next generation saRNA platform for RNA therapeutics. Combining these optimized saRNA vectors with advanced delivery technologies, such as ligand conjugated targeted LNP, may further enhance the precision, efficacy, and therapeutic potential of mRNA-based drugs.

## Results

### 1 Comprehensive optimization the design of saRNA based on VEEV

Although the VEEV based saRNA vector is widely used in mRNA drug development, further optimization is needed to enhance protein expression, while reducing immunogenicity. In this study, we optimized the VEEV saRNA vector using an EGFP-expressing construct, systematically testing different UTRs of subgenomic RNA, co-factors, capping analogs, and poly(A) tails to improve its performance.

In conventional saRNA design, the target gene ORF is typically placed directly after the subgenomic (SG) promoter. Since UTRs enhance protein expression in linear mRNA by recruiting ribosomes through their unique secondary structures^30^, adding a 5’ UTR after the SG promoter could theoretically boost subgenomic RNA translation. To test this, we inserted four different 5’ UTRs at the 5’ end of the EGFP sequence and compared the expression of EGFP saRNAs with and without the SG-5’ UTR in HEK293T cells. At 24- and 48-hours post-transfection, EGFP saRNAs containing the SG-5’ UTR showed a 1.8-fold increase in expression compared to those without it (Fig. 1B). Among the tested UTRs, UTR-1 exhibited the highest enhancement in expression and was selected for further optimization. These results confirm that adding an SG-5’ UTR enhances target protein expression in saRNA.

**Figure 1.**
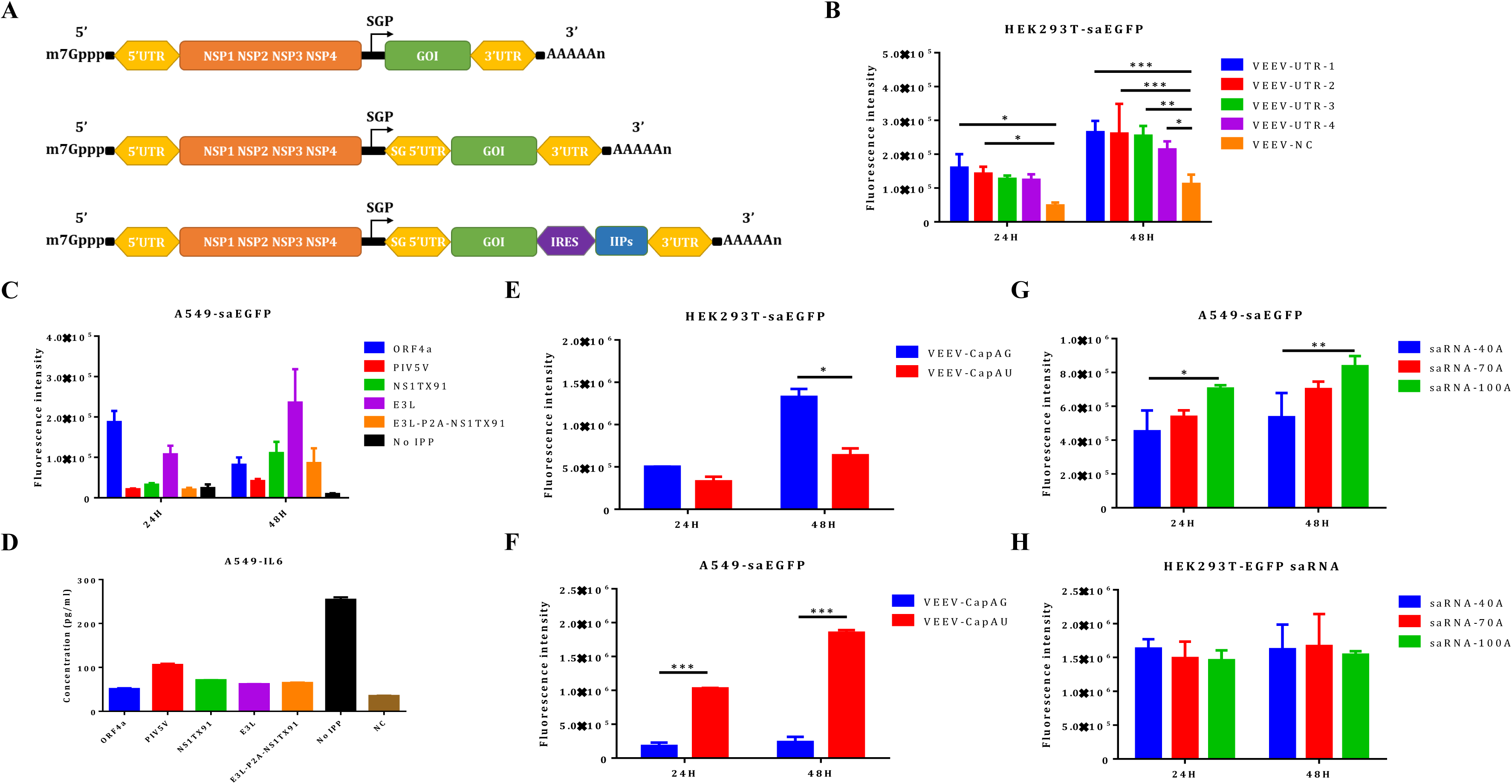
Comprehensive optimization of VEEV-based saRNA constructs. (A) Schematic illustration of the optimization process, including incorporation of a subgenomic 5’ UTR, co-expression of immunosuppressive proteins via CVB3 IRES, and evaluation of capping structures and polyA tail lengths. (B) EGFP expression levels in HEK293T cells transfected with 10ng saRNA constructs containing various subgenomic 5’ UTRs using Lipofectamine. (C-D) EGFP expression (C) and IL-6 levels (D) in A549 cells following transfection with 10ng EGFP saRNA co-expressing different immunosuppressive proteins using Lipofectamine. (E-F) EGFP expression in HEK293T (E) and A549 (F) cells transfected with 10ng EGFP saRNA containing different cap structures. (G-H) EGFP expression in HEK293T (G) and A549 (H) cells transfected with 10ng EGFP saRNA containing different polyA tail lengths. Data shown as mean + SD of 3 independent experiments.

During replication, saRNA forms dsRNA, which can trigger innate immune activation through pathways like RIG-I/MDA5, PKR, and others^1^. This immune response can lead to RNA degradation, translation inhibition, and reduced protein expression. To enhance saRNA expression stability, modulating immune activation is a crucial strategy. Previous studies show that co-transfecting immunosuppressive proteins (IPPs) or co-expressing them via 2A peptides can significantly improve saRNA expression by reducing immune activation^21,31^. We selected four immunosuppressive proteins (ORF4a, PIV5V, NS1TX91, and E3L) and co-expressed them in saRNA using the CVB3 IRES to evaluate their effectiveness. IRES was chosen over 2A peptides to avoid potential interference from 2A sequences and to maintain lower translation levels, reducing the risk of excessive IPP expression disrupting normal cellular functions. The results showed that 48 hours post-transfection, all IPP co-expression groups significantly increased EGFP expression by 4- to 23-fold (Fig. 1C), and reduced the immune-related cytokine IL-6 by more than 2.5-fold (Fig. 1D). Since E3L and NS1 TX91 showed the highest expression enhancement, we co-expressed them using P2A to determine if combining multiple IPPs would further improve performance. However, P2A co-expression resulted in lower expression levels than individual IPPs, possibly due to the 2A peptide sequence affecting translation.

The positive-sense strand genomes of alphaviruses typically begin with a 5′ AU structure. For saRNA capping, using the AU cap is recommended because it preserves the authentic alphavirus 5′ end, leading to efficient capping and higher yield^15^. However, the impact of other cap analogs on the performance of saRNA is not yet well understood. Here, we wanted to test whether changing AU to AG would affect the expression of the saRNA. We designed two different T7 transcription initiation sequences, TATA<u>AT</u> and TATA<u>AG</u>AT, for efficient Cap-AG and AU co-transcriptional capping of EGFP saRNA. The expression of these EGFP saRNA were tested in HEK293T cells and A549 cells. The results showed that in HEK293T cells, AG-capped saRNA expression was approximately 1.9 times higher than that of AU-capped saRNA at 48 hours post-transfection (Fig. 1E). However, in A549 cells, AG-capped saRNA expression was only about 13% of the AU-capped saRNA expression at 48 hours post-transfection (Fig. 1F). Considering that A549 cells exhibit immune responses more similar to the in vivo environment compared to HEK293T cells, we recommend using Cap-AU for saRNA capping.

Finally, we tested the effect of polyA tail length on saRNA expression. Longer polyA tails in conventional mRNA are known to improve stability and translation. We compared saRNAs with polyA tails of 40A, 70A, and 100A in both HEK293T and A549 cells. In A549 cells, 100A polyA tails enhanced expression by 1.6 times compared to 40A, but no significant difference was observed between 70A and 100A (Fig. 1G). In HEK293T cells, no differences were noted across the polyA tail lengths (Fig. 1H). Based on these results, a 70A polyA tail was selected for further optimization.

In conclusion, our findings suggest that a combination of UTR optimization, immune suppression, and careful selection of capping and polyA strategies can significantly improve the VEEV based saRNA expression, and this strategy can be extended to other alphavirus based saRNA vector constructs.

### 2 In vitro screening for alphavirus vectors with enhanced protein expression

In addition to VEEV, the alphavirus genus comprises over 30 subgroups, including well-studied members like Semliki Forest virus (SFV), Sindbis virus (SINV), and Eastern equine encephalitis virus (EEEV), which can largely be categorized into seven distinct complexes (Supplementary Figure 2). To evaluate whether saRNA derived from other alphaviruses could achieve better protein expression than VEEV, 28 alphavirus family members were selected and engineered to express EGFP for testing. These EGFP saRNAs were tested for expression in HEK293T and A549 cells, using the original VEEV-based EGFP saRNA as a control. In addition, saRNA constructs with and without m5C modifications were compared, as previous studies have shown that m5C modifications enhance the expression of VEEV-based saRNA. However, whether this effect extends to saRNAs derived from other alphavirus subtypes remains unclear.

The results showed that at 48 hours post-transfection, 7 out of the 28 tested EGFP saRNAs derived from different alphaviruses exhibited more than four times the EGFP expression level of the control VEEV EGFP saRNA in both HEK293T and A549 cells (Fig. 2A-B, Supplementary Figure 3). These included Everglades virus (EVEV), Highlands J virus (HJV), Mosso das Pedras virus (MDPV), Mucambo virus (MUCV), Ndumu virus (NDUV), Pixuna virus (PIXV), and Rio Negro virus (RNV). Additionally, in both HEK293T and A549 cells, m5C modifications significantly enhanced the expression of HJV, Sagiyama virus (SAGV), and VEEV saRNAs compared to their unmodified counterparts. In contrast, m5C modifications led to a significant decrease in expression for the NDUV and Trocara virus (TROV) groups. For the remaining saRNA constructs, the effects of m5C modification varied between cell types or showed no significant differences.

**Figure 2.**
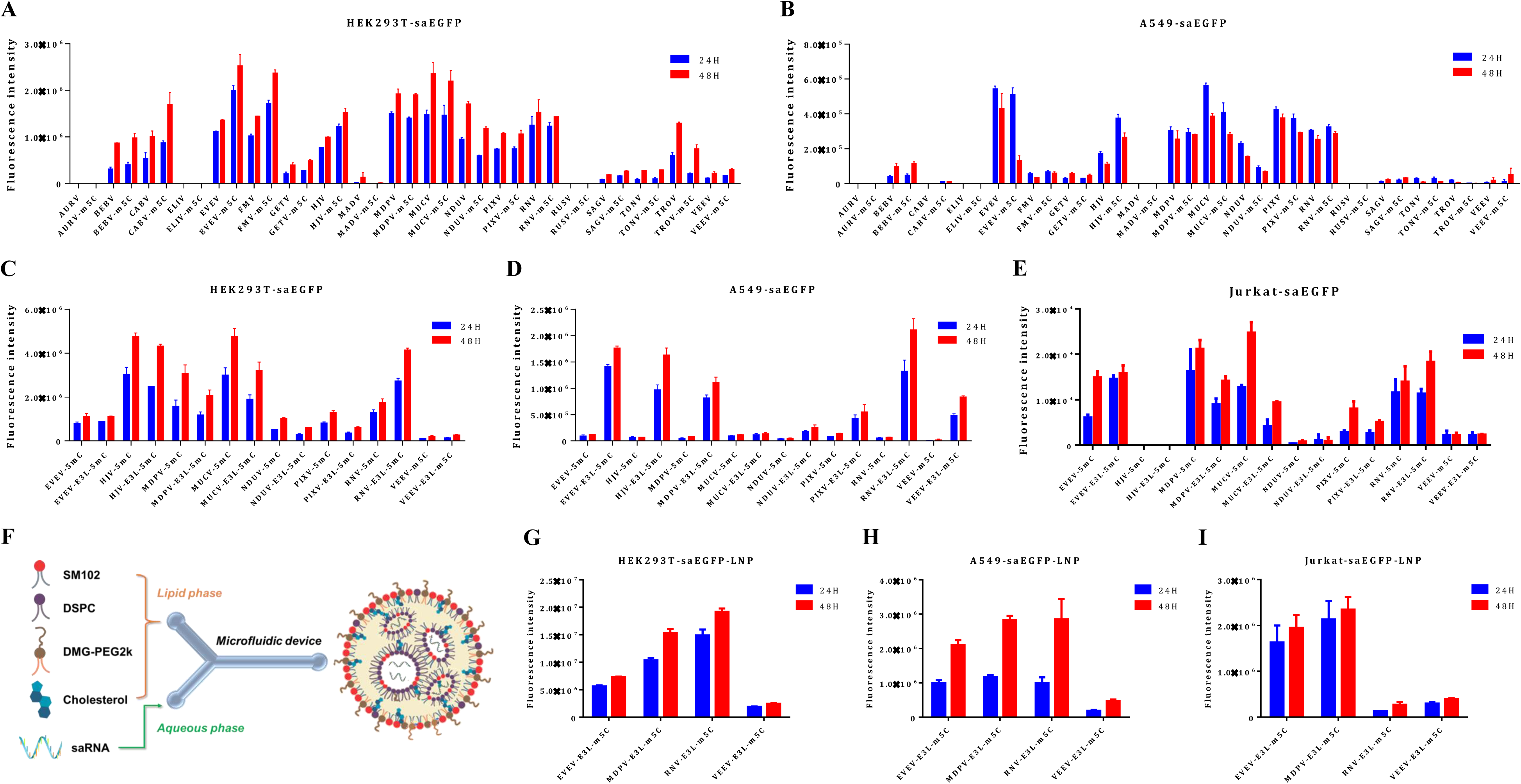
Systematic in vitro screening of alphavirus-derived saRNAs. (A-B) EGFP expression in HEK293T (A) and A549 cells (B) following transfection with 10ng unmodified or m5C-modified EGFP saRNAs derived from various alphaviruses using Lipofectamine. (C-E) EGFP expression in HEK293T (C), A549 (D), and Jurkat cells (E) after transfection with 10ng EGFP saRNAs, with or without co-expression of the immunosuppressive protein E3L, using Lipofectamine. (F) Schematic illustration of the encapsulation process for saRNA-loaded lipid nanoparticles (LNPs). (G-I) EGFP expression in HEK293T (G), A549 (H), and Jurkat cells (I) transfected with 10ng EVEV-, MDPV-, RNV-, and VEEV-derived EGFP saRNAs formulated in SM102 LNPs. Data shown as mean + SD of 3 independent experiments.

Notably, the EGFP fluorescence intensity of the different saRNA groups in A549 cells was significantly lower than in HEK293T cells, potentially due to the immunogenicity of self-amplifying RNA. To investigate whether immunogenicity reduction could enhance expression, co-expression of the E3L protein with these saRNAs using CVB3 IRES was examined. The expression of EGFP-CVB3-E3L saRNAs derived from EVEV, HJV, MDPV, MUCV, NDUV, PIXV, and RNV was evaluated in HEK293T, A549, and Jurkat cells. The results indicated that, similar to VEEV, co-expression of E3L enhanced EGFP expression by more than fourfold in A549 cells for all saRNA groups, except for the MUCV group (Figure. 2D). However, in HEK293T and Jurkat cells (Figure. 2C, 2E), the inclusion of the E3L sequence did not lead to increased expression.

To assess the efficiency of LNP delivery for saRNA, the expression of SM102-based LNP-encapsulated saRNA was evaluated in HEK293T, A549, and Jurkat cells. Along with the control VEEV, three novel saRNA constructs—EVEV, DMPV, and RNV— exhibiting strong expression in all three cell lines were encapsulated in LNP formulated with SM102 and tested. Robust expression was observed across all tested saRNAs (Figures. 2F-H). In all three cell lines, EGFP expression levels of EVEV- and MDPV-derived saRNA exceeded those of the VEEV group by more than threefold. Notably, the highest expression levels were achieved by RNV-derived EGFP saRNA-LNP in HEK293T and A549 cells, surpassing the VEEV group by over sixfold. However, in Jurkat cells, its expression was the lowest, significantly reduced compared to the VEEV group. These findings suggest that replication efficiency and compatibility of different alphavirus-derived saRNAs may vary depending on cell type, potentially reflecting differences in their replication mechanisms or interactions with the cellular environment.

### 3 In Vivo Expression of Different saRNAs

To evaluate the duration and magnitude of expression of different saRNAs in vivo, VEEV-, MDPV-, RNV-, and EVEV-derived saRNAs were constructed and synthesized, along with a linear mRNA control encoding Firefly Luciferase (FLuc). The FLuc saRNAs and mRNA were encapsulated using an SM102-based LNP formulation. The physicochemical properties of each LNP are summarized in Supplementary Table 1. Both FLuc saRNA-LNP and FLuc mRNA-LNP exhibited a polydispersity index (PDI) of less than 0.2, indicating good dispersion and uniformity of the LNP particles. Zeta potential measurements ranged between ±15 mV, suggesting favorable electrostatic neutrality in a neutral buffer environment. Additionally, the encapsulation efficiency of all RNA-LNP formulations exceeded 92%. Further characterization of FLuc mRNA-LNP and VEEV-derived FLuc saRNA-LNP using cryo-electron microscopy (cryo-EM) revealed uniform particle size and well-defined morphology (Supplementary Figure 4).

Following encapsulation, these LNP formulations were administered to mice via tail vein injection at a dose of 0.25 mg/kg. Bioluminescence imaging was performed on days 1, 2, 3, 6, 10, 14, and 21 to monitor FLuc expression over time. Besides, mouse body weight (Figure. 3B) was monitored to evaluate safety. No significant differences in weight were observed across all treatment groups, including the control group, suggesting good tolerability of LNP-delivered saRNAs. The Bioluminescence results indicated distinct expression patterns between FLuc mRNA and FLuc saRNA in vivo (Figure. 3C-D). FLuc mRNA exhibited the highest luminescent signals on day 1, followed by a rapid decline, reaching levels comparable to the PBS control group by day 10. In contrast, the luminescent signals of FLuc saRNA were over 40-fold lower than those of FLuc mRNA on day 1. However, FLuc saRNA expression remained stable across all groups until day 6, after which a gradual decline was observed. Notably, even at day 21, luminescent signals in the FLuc saRNA groups remained at a relatively high level, exceeding those of the FLuc mRNA group by more than threefold.

**Figure 3:**
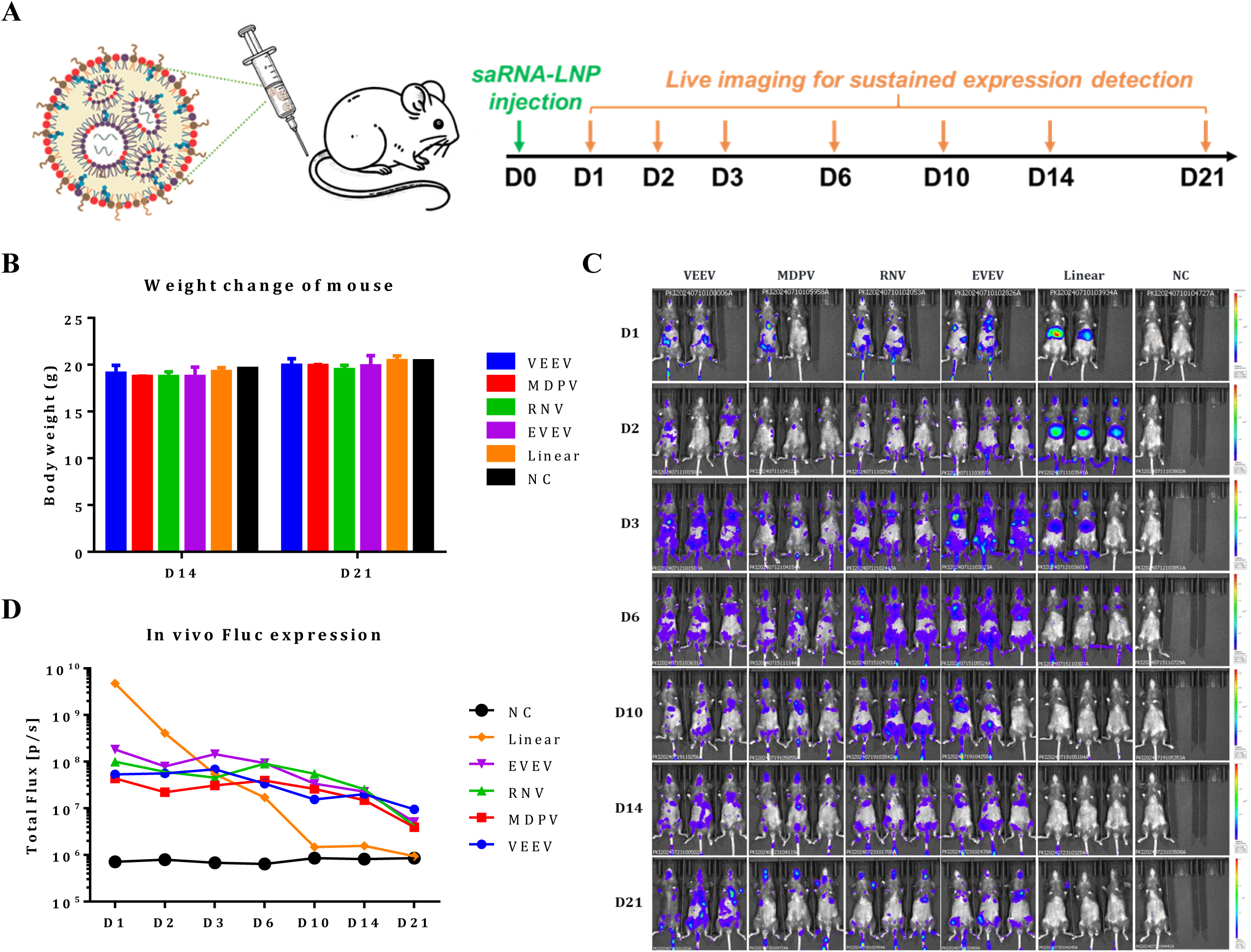
In vivo test of mice injected with FLuc saRNA-LNPs or mRNA-LNP. (A) Schematic illustration of the timeline for mouse injections and subsequent detection procedures. (B) Weight change of mouse injected with 5 μg FLuc saRNA or mRNA formulated with SM102 LNP via intramuscular (IM) route of administration. (C-D) Representative images and luminescence intensity of mice injected with 5 μg of FLuc saRNA or mRNA LNP. Luminescence was measured and quantified using an In Vivo Imaging System (PerkinElmer).

### 4 Organ specific expression of Fluc-saRNAs

To investigate the tissue-specific distribution of saRNA expression, bioluminescence signals were analyzed in various organs, including the heart, liver, spleen, lung, kidney, brain, and lymph nodes, 24 hours after administration. We also aimed to test whether the in vivo distribution of saRNA-LNP is different from that of mRNA-LNP. Since saRNA is longer than conventional mRNA, its packaging into LNPs may exhibit distinct delivery characteristics. To investigate this, DiR fluorescent dye was incorporated into LNPs encapsulating both saRNA and mRNA to analyze their distribution. Distinct expression patterns were observed between FLuc mRNA and saRNA in vivo (Figure. 4A-C). As reported previously, FLuc expression in the FLuc mRNA-LNP group was mainly concentrated in the liver, representing 93.53% of the total luminescent signals, with a smaller fraction observed in the spleen (3.16%). In contrast, FLuc saRNA-LNP groups showed considerable extrahepatic expression, with liver-derived FLuc signals making up only 2.07%–8.23% of the total. Remarkably, distinct tissue tropisms were noted across different FLuc saRNA groups. In the VEEV group, FLuc expression was distributed among the kidney, spleen, and heart, each contributing over 20% of the total signal, with the highest proportion seen in the spleen (29.24%). For the MDPV group, FLuc expression was predominantly found in the lymph nodes (47.14%) and spleen (38.72%). The RNV and EVEV groups demonstrated the highest expression in the spleen, accounting for 52.83% and 72.68%, respectively. Additionally, FLuc expression in the brain was minimal across all groups, with levels below 2.5%. Regarding absolute luminescent signal intensity, the FLuc mRNA-LNP group showed the highest expression in the liver, while the VEEV group displayed the strongest expression in non-hepatic organs. The EVEV group also exhibited strong spleen-specific luminescent signals, comparable to the VEEV group, both significantly higher than those seen in the FLuc mRNA-LNP group. These findings highlight the unique tissue tropism of saRNAs compared to linear mRNA.

**Figure 4.**
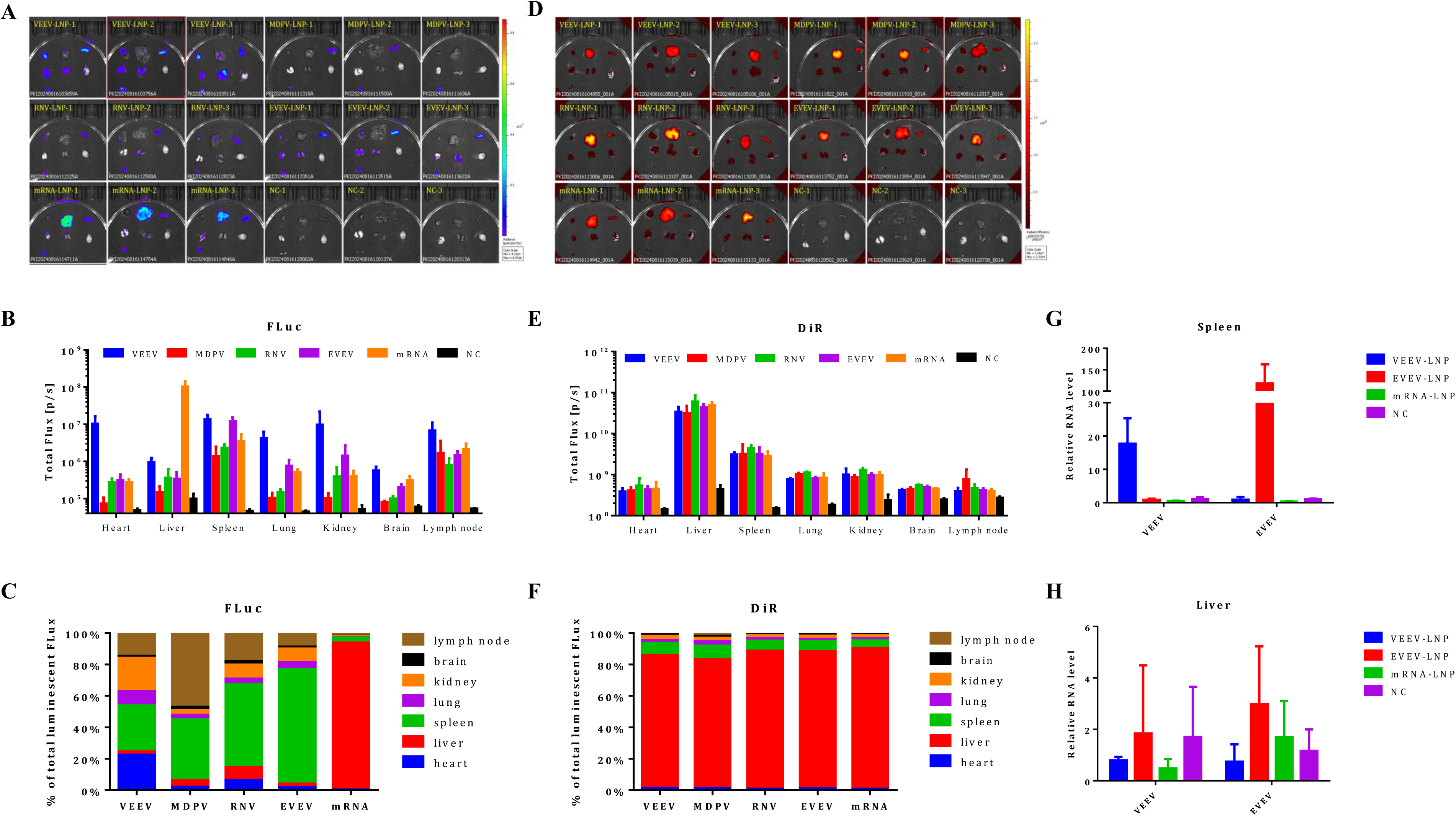
In vivo expression and biodistribution of FLuc saRNA-LNPs and mRNA-LNP in mice. (A–C) Representative bioluminescence images (A), signal intensity (B), and organ-specific distribution ratios (C) in mice intravenously injected with 5 μg of FLuc saRNA-LNP or mRNA-LNP. (D–F) Representative DiR fluorescence images (D), signal intensity (E), and organ distribution ratios (F) showing biodistribution of FLuc saRNA-LNP or mRNA-LNP 24 hours post-injection. (G–H) RNA levels of VEEV and EVEV saRNAs in the spleen (G) and liver (H) of injected mice, quantified by RT-qPCR using the 7500 Fast Real-Time PCR System (Thermo Fisher Scientific, USA). Luminescence was measured using the In Vivo Imaging System (PerkinElmer).

For LNP distribution, the results (Figure. 4D-F) showed that 24 hours after injection of RNA-LNP, DiR signals from both FLuc saRNA-LNP and FLuc mRNA-LNP were primarily localized in the liver (over 82%), with a smaller proportion in the spleen (5%-9%), indicating that the SM102-based LNP primarily delivers RNA to the liver. This suggests that the longer length of saRNA does not significantly alter the overall distribution of LNPs compared to conventional mRNA-LNPs, and the observed extrahepatic expression of saRNA is not influenced by its delivery distribution.

Based on the results above and previous reports^32^, it was hypothesized that the replication efficiency of saRNA varies across organs, leading to tissue-specific expression. To test this, RT-qPCR was conducted to measure the levels of non-structural protein RNA in the liver and spleen following administration of VEEV-LNP and EVEV-LNP. As shown in Figure. 4G-H, both the VEEV-LNP and EVEV-LNP groups exhibited over a 10-fold higher level of non-structural protein RNA in the spleen compared to the liver. This demonstrates that saRNA can efficiently replicate in the spleen. Although a relatively small proportion of RNA is delivered to the spleen, efficient replication enables high-level protein expression. These findings highlight the critical role of replication efficiency in determining saRNA’s tissue-specific expression and provide insights for optimizing saRNA-based applications to achieve targeted expression in extrahepatic tissues.

## Discussion

VEEV-based saRNA vector is widely utilized for RNA therapeutics research, with ongoing optimization focusing on mutant non-structural proteins and co-expression of immunosuppressive proteins to enhance performance^9,19,33,34^. In this study, we systematically optimized the saRNA design by evaluating critical components such as cap structures, polyA tail lengths, subgenomic 5’ UTR sequences, m5C modifications, and immunosuppressive protein (IPP) candidates. Through comprehensive evaluation, we identified an optimized VEEV saRNA design featuring Cap-AU capping, a subgenomic 5’ UTR, co-expression of the IPP E3L via CVB3 IRES downstream of the target gene, and a 70-nucleotide polyA tail. This optimized vector substantially increased target gene expression. We further expanded our analysis by engineering 28 saRNA vectors derived from different alphaviruses and assessed their expression in vitro. The top three ranked saRNA vectors (EVEV, MDPV, and RNV) were subsequently evaluated in vivo for their tissue-specific expression profiles using the luciferase reporter. For the first time, our results demonstrated that 1) the optimized VEEV vector exhibited the highest expression across various non-liver tissues; 2) EVEV saRNA primarily localized to the spleen, achieving expression levels comparable to VEEV; 3) MDPV and RNV saRNAs predominantly expressed in spleen and lymph nodes but at lower intensities, with MDPV favoring lymph nodes and RNV showing higher spleen expression. These findings highlight superior candidates with distinct tissue preferences for targeted RNA delivery.

Given the structural similarity between saRNA-derived subgenomic RNAs and conventional mRNAs, we anticipated that incorporating 5’ UTRs would enhance saRNA protein expression^35^. Indeed, adding these UTR sequences significantly boosted saRNA target protein expression. Unlike linear mRNA, where longer polyA tails generally enhance stability and translation^36,37^, saRNA expression was less sensitive to polyA tail length changes. This may be due to the polyA tail primarily influencing non-structural protein translation, whereas subgenomic RNA independently acquires its polyA tail after replication. Previous reports indicate that common uridine modifications [e,g, pseudouridine (ψ), N1-methylpseudouridine (m1ψ)] effective in conventional mRNA, can significantly reduce saRNA expression, whereas cytidine modifications, specifically 5-methylcytidine (m5C) and hydroxymethylcytidine (hm5C), notably enhance VEEV saRNA expression^38,39^. Consistent with these reports, our data also confirmed this conclusion (Supplementary Figure. 1). This may be due to uridine modifications altering saRNA structural integrity, affecting replication efficiency, given saRNA’s replication reliance on replicon recognition of conserved 5’ and 3’ sequences and subgenomic promoters. Interestingly, replacing the conventional AU cap with an AG cap significantly enhanced saRNA expression in HEK293T cells but markedly reduced expression in immune-responsive A549 cells. This suggests that optimization strategies improving non-structural protein expression might simultaneously increase immunogenicity, triggering innate immune responses and suppressing target protein translation. This pattern was consistently observed during alphavirus vector screening. Co-expression of the immunosuppressive protein E3L mitigated immunogenicity for some saRNAs, significantly improving expression. However, other saRNAs remained ineffective despite E3L co-expression, indicating that non-structural proteins may inherently differ in their immune activation potential, probably through differential engagement of host pattern recognition receptors (such as RIG-I, MDA5, or PKR)^24,26^. Consequently, careful assessment of immunogenic profiles should be integral to the optimization process for saRNA vectors, alongside strategies aimed at enhancing protein expression and replication efficiency.

Extrahepatic targeting of mRNA delivery has been a key research focus in the field of mRNA therapeutics^40–42^. In this study, we were surprised to find that intravenously injected FLuc saRNA-LNP primarily exhibited expression outside the liver in mice, even though tracking the distribution of DiR-labeled LNPs revealed that, after injection, saRNA-LNPs predominantly accumulated in the liver. Moreover, saRNAs derived from different alphaviruses showed distinct preferences for accumulation in extrahepatic organs. Alphaviruses are classified into Old World (OW) and New World (NW) groups based on their geographical origins and pathogenic profiles^23^. OW alphaviruses, such as Chikungunya virus (CHIKV) and Ross River virus (RRV), primarily targeting musculoskeletal tissues like muscle satellite cells and osteoblasts, leading to myositis and arthritis^43,44^. In contrast, NW alphaviruses, including Eastern equine encephalitis virus (EEEV) and VEEV, exhibit neurotropic tendencies, infecting neurons and glial cells in the central nervous system, resulting in encephalitis^45,46^. Additionally, alphaviruses demonstrate a strong tropism for immune organs such as the spleen and lymph nodes, where they replicate efficiently in macrophages and dendritic cells, triggering robust innate immune responses^25,26^. This led us to hypothesize that the extrahepatic expression of saRNA might be related to the replication characteristics of alphaviruses replicon. RT-qPCR analysis of different tissues further confirmed this assumption, showing high levels of saRNA in the spleen but very low levels in the liver. This result is consistent with recent findings reported by Nuthan Vikas Bathula et al^32^. They compared the distribution and expression dynamics of saRNA delivered via LNP or pABOL in mice under different injection routes. Their findings revealed that, regardless of the administration method, saRNA delivered by LNP or pABOL exhibited extrahepatic replication and expression characteristics in mice. Therefore, the tissue-specific expression patterns of saRNA likely result from varying replication efficiencies influenced by local immune responses and host factors. Future optimization efforts could explore manipulating these interactions to further refine and enhance saRNA’s targeted expression profiles in specific tissues.

Among the tested saRNAs, VEEV and EVEV demonstrated the highest in vivo expression and notably distinct tissue distribution profiles. VEEV-derived saRNA had extensive extrahepatic expression, with strong protein expression in the spleen and heart, which differs from previous findings, where VEEV infection was mainly observed in the brain, spleen, and lungs of captured fruit bats^47^. The divergent tropism observed in this study likely results from the VEEV-TC83 strain used, which underwent 83 serial passages in guinea pig heart cell cultures, potentially altering its genomic characteristics and shifting its tissue specificity toward cardiac and broader extrahepatic expression^48^. The enhanced cardiac expression of VEEV-derived saRNA highlights its potential as a novel approach for heart-targeted therapies, particularly given that conventional mRNA-LNP formulations exhibit limited cardiac targeting efficiency. Additionally, continuously passaging alphavirus-derived saRNA through specific cell lines may serve as an effective strategy to evolve and enhance their tissue-specific targeting capabilities. In contrast, EVEV-derived saRNA exhibited remarkable spleen specificity, with 72.68% of its FLuc signal concentrated in the spleen, far higher than other saRNA groups. It can significantly reduce potential liver toxicity while improving targeted tissue specificity, positioning it as a highly promising candidate to enhance the effectiveness of immunotherapies^49^, such as gene editing in immune cells and the optimization of CAR-T cell therapies. Furthermore, due to its focused immunological targeting and reduced systemic toxicity, it holds particular promise for vaccine development, especially for conditions where robust antigen-specific immune responses are desired. We have been successfully utilized saRNA-LNP immunization for antibody discovery against difficult targets such as multi-transmembrane proteins like GPCRs in both mice and rabbits. Unexpectedly, it outperformed conventional mRNA-LNP, plasmid DNA, and protein-based immunizations (hCCR9, unpublished data). Moving forward, testing these saRNA vectors in non-human primates would help validate their translational potential and organ-targeting specificity in clinically relevant models.

Our study conclusively demonstrates that optimized saRNA expression can be achieved through the addition of 5’ UTRs, co-expression of immune suppression proteins, and testing various transcription start sites and polyA tail lengths. For the first time, we identified multiple alphavirus-derived saRNA backbones with superior expression, particularly in immune-sensitive cells, and demonstrated that LNP delivery significantly enhanced saRNA expression across different cell types. Our in vivo studies further revealed that saRNAs exhibit sustained expression and broader tissue distribution compared to linear mRNA, with tissue-specific expression being largely driven by replication efficiency. These findings provide important insights for optimizing saRNA-based therapies and highlight the need to consider both viral backbone selection and delivery systems in the design of effective saRNA-based vaccines and gene therapies.

## MATERIALS AND METHODS

### Plasmids and strains

The saRNA was designed by replacing the structural protein sequence of the VEEV (TC-83 strain) with the EGFP protein encoding gene. The construct includes a mini T7 promoter at the 5’ end and a 70bp poly A tail with a BspQI cleavage site at the 3’ end. For vectors containing sub-genomic 5’ UTRs, different UTR sequences were inserted between the sub-genomic promoter and the EGFP sequence. Vectors designed to co-express immunosuppressive proteins had these proteins linked to the EGFP sequence using the CVB3 IRES. For vectors employing various capping methods, the transcription start sequence following the T7 promoter TATA-box was modified to the specified sequence. The saRNAs for Alphavirus screening were developed by replacing the structural protein sequences of various Alphaviruses with the EGFP gene. Those derived from EVEV, HJV, MDPV, MUCV, NDUV, PIXV, and RNV were further engineered to include a sub-genomic 5’ UTR and co-express immunosuppressive proteins via an IRES sequence. Additionally, saRNAs encoding FLuc were constructed by substituting the EGFP sequence with the firefly luciferase gene. For the conventional FLuc mRNA, the encoded firefly luciferase protein was flanked by the 5’ and 3’ UTRs of the human α-globin gene and featured a mini T7 promoter at the 5’ end along with a 70bp poly A tail and a BspQI cleavage site at the 3’ end. All sequence information is detailed in Supplementary file 1.

### IVT mRNA synthesis

Template DNA for IVT was generated by linearization of the vectors using the BspQI restriction enzymes (NEB). After linearization, the IVT reactions to generate mRNA and saRNA were carried out using different methods. Conventional mRNA was co-transcription capped with a cap-AG analogue (Syngenebio) and modified by replacing UTP with N1-methylpseudo-UTP (m1Ψ) in the transcription reaction. The saRNA was co-transcription capped with a cap-AU analogue (Syngenebio) and unmodified or modified by replacing CTP with 5-methyl-CTP (m5C) in the transcription reaction. The mRNA and saRNAs were purified by lithium chloride precipitation and dissolved in RNase-free water. For animal study, the mRNA and saRNAs were further purified using HPLC with an oligo dT column. The integrity of the RNA was assessed by capillary electrophoresis using 5200 Fragment Analyzer system (Agilent).

### Cell culture and RNA transfection

The human embryonic kidney cell line HEK293T, human non-small cell lung cancer cell line A549, and immortalized human T lymphocyte cell line Jurkat were obtained from the American Tissue Culture Collection. HEK293T and A549 cells were grown in high-glucose Dulbecco’s Modified Eagle Medium (DMEM) supplemented with 10% fetal bovine serum (FBS, Gibco) and 1% penicillin-streptomycin (Gibco). Jurkat cells were maintained in RPMI-1640 medium with 10% FBS (Gibco) and 1% penicillin-streptomycin (Gibco). All cells were incubated at 37 °C with 5% CO2. RNAs were transfected into the cells using Lipofectamine MessengerMAX Reagent (Thermo), following the manufacturer’s protocol. For LNP delivery, RNA-LNP complexes were directly added to the cultured cells. The EGFP signal was detected and imaged by fluorescence microscopy, and quantified by Image J software. Firefly Luciferase activity was measured in an Infinite 200 PRO microplate reader (TECAN) using the Luciferase Assay System (Promega). IL-6 protein levels were quantified from supernatants by ELISA (R&D Systems).

### Lipid nanoparticle formulation and characterization

The saRNAs and mRNAs were encapsulated into lipid nanoparticles (LNPs) using an SM102-based formulation. Specifically, 1,2-distearoyl-sn-glycero-3-phosphocholine (DSPC), SM102, cholesterol, and 1,2-dimyristoyl-rac-glycero-3-methoxypolyethylene glycol-2000 (DMG-PEG-2000) were sourced from Sinopeg (CN). A lipid mixture was prepared in ethanol, consisting of SM102, DSPC, cholesterol, and DMG-PEG-2000 at a molar ratio of 50:10:38.5:1.5. For formulations labeled with DiR fluorescent dye, DiR was added into the lipid mixture for LNP fluorescent labeling at 0.5% molar rate, maintaining the same lipid ratio, to achieve fluorescently traceable LNPs. The RNA aqueous phase was prepared by diluting saRNA or mRNA transcripts in 25 mM sodium acetate buffer (pH 4.53). Using a Microflow S Microfluidic device (Mingtai), the lipid and aqueous phases were rapidly mixed at an N:P ratio of 6, with a total flow rate of 12 mL/min and a 1:3 lipid-to-RNA flow rate ratio. The resulting LNP formulations were diluted 5× with PBS and concentrated via buffer exchange using Amicon filters (100K MWCO). Encapsulation efficiency of the saRNA or mRNA in LNPs was assessed using the Quant-iT Ribogreen RNA assay kit (Invitrogen, Thermo Fisher Scientific). Particle size, polydispersity index (PDI), and zeta potential were measured via dynamic light scattering (DLS) using the Zetasizer Pro (Malvern Instruments).

### Mouse and in vivo administration

All national and institutional guidelines were strictly followed for the housing and care of mice at Cyagen Biosciences (CN). Animal experiments were performed in accordance with protocols approved by the Cyagen Biosciences Group Institutional Animal Care and Use Committee (IACUC). Female BALB/c mice, 8 weeks old, were housed in groups of 3–5 per cage in a fully acclimatized environment with sterile food and water provided ad libitum. Each mouse received a 100 μL intravenous injection containing 5 μg of FLuc mRNA-LNP, FLuc saRNA-LNP, or PBS.

### In vivo imaging

On the imaging day, mice were intraperitoneally injected with 100 μL of 30 mg/mL D-luciferin Bioluminescent Substrate (PerkinElmer, US). After the injection, the mice rested for 7 minutes before being anesthetized with 2–3% isoflurane for 3 minutes. Imaging was performed approximately 10 minutes after the D-luciferin injection using the IVIS Spectrum In Vivo Imaging System (PerkinElmer). The bioluminescent signals were quantified post-imaging using Living Image® 4.5.5 Software (IVIS Imaging Systems) and expressed as total flux (photons/second).

### Real time quantitative PCR

For quantitative PCR (qPCR) analysis, total RNA was extracted from mouse tissues using the FastPure Complex Tissue/Cell Total RNA Isolation Kit (Vazyme, CN) following the manufacturer’s protocol. The RNA concentration was measured using NanoDrop, and reverse transcription into cDNA was performed using the HiScript III RT SuperMix (Vazyme, CN). Primer sequences specific to mouse GAPDH and the nonstructural proteins of VEEV or EVEV are listed in Supplementary file 1.

qPCR was carried out using the ChamQ Universal SYBR qPCR Master Mix (Vazyme, CN) according to the manufacturer’s instructions. The reaction mix was prepared on ice, including cDNA template, 1×master mix with ROX reference dye and 0.2 μM of each primer. The qPCR was performed using the 7500 Fast Real-Time PCR system (Thermo Fisher Scientific, USA). The protocol included a 30-second pre-incubation at 95 for polymerase activation, followed by 40 cycles of amplification: 10 seconds at 95 for denaturation and 30 seconds at 60 for extension. The relative levels of saRNA transcripts in total RNA were calculated using the ΔΔCt method.

### Statistics

All data are the result of three independent experiments, unless declared otherwise, and are represented as mean ± standard deviation. Experiments were analyzed for statistical significance with one-way ANOVA followed by the Bonferroni post hoc test. Statistical analysis was performed using GraphPad Prism 6 software. Asterisks are used to illustrate statistical significance (* p < 0.05, ** p< 0.005, *** p < 0.0001).

---

**Supplementary Figure 1.**
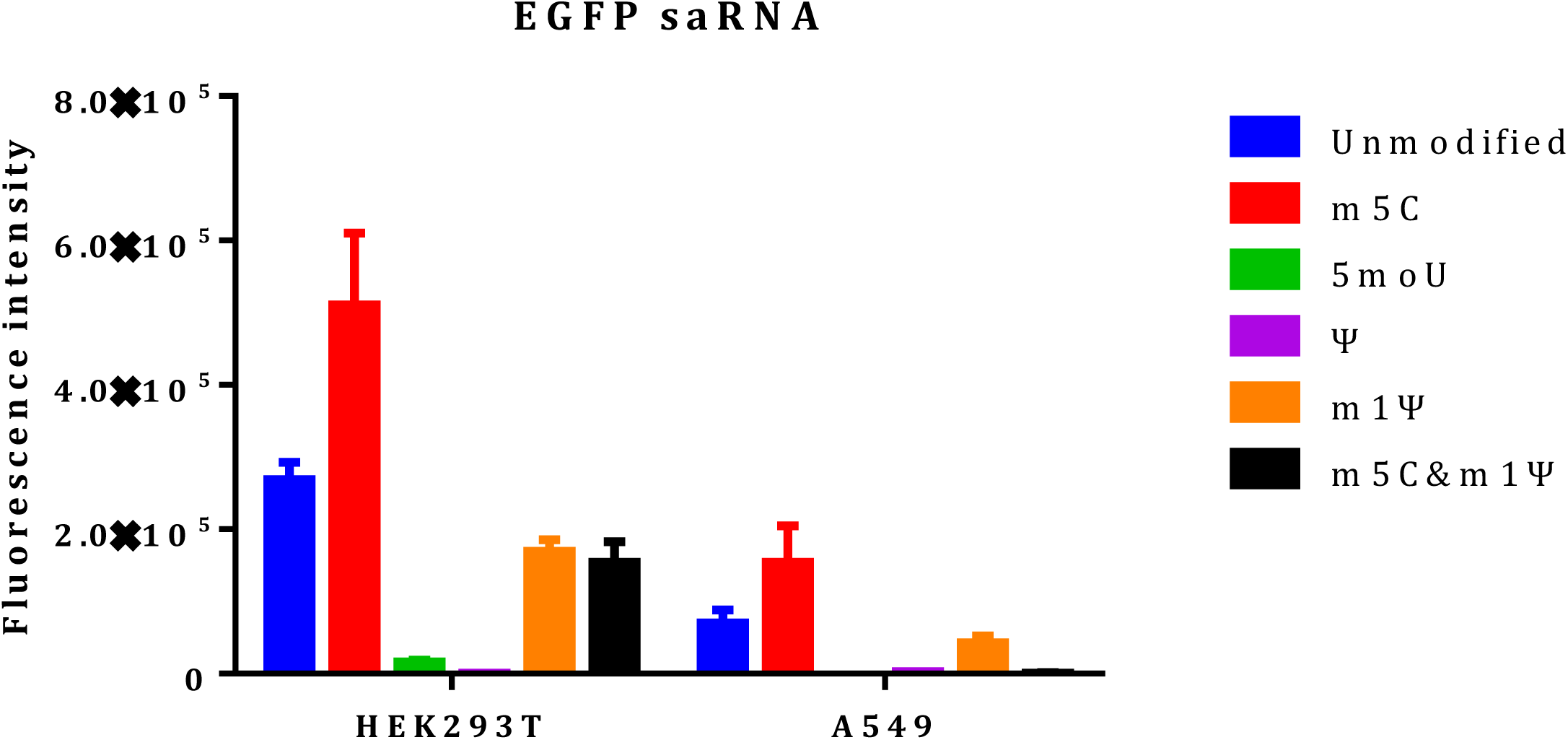

**Supplementary Figure 2.**
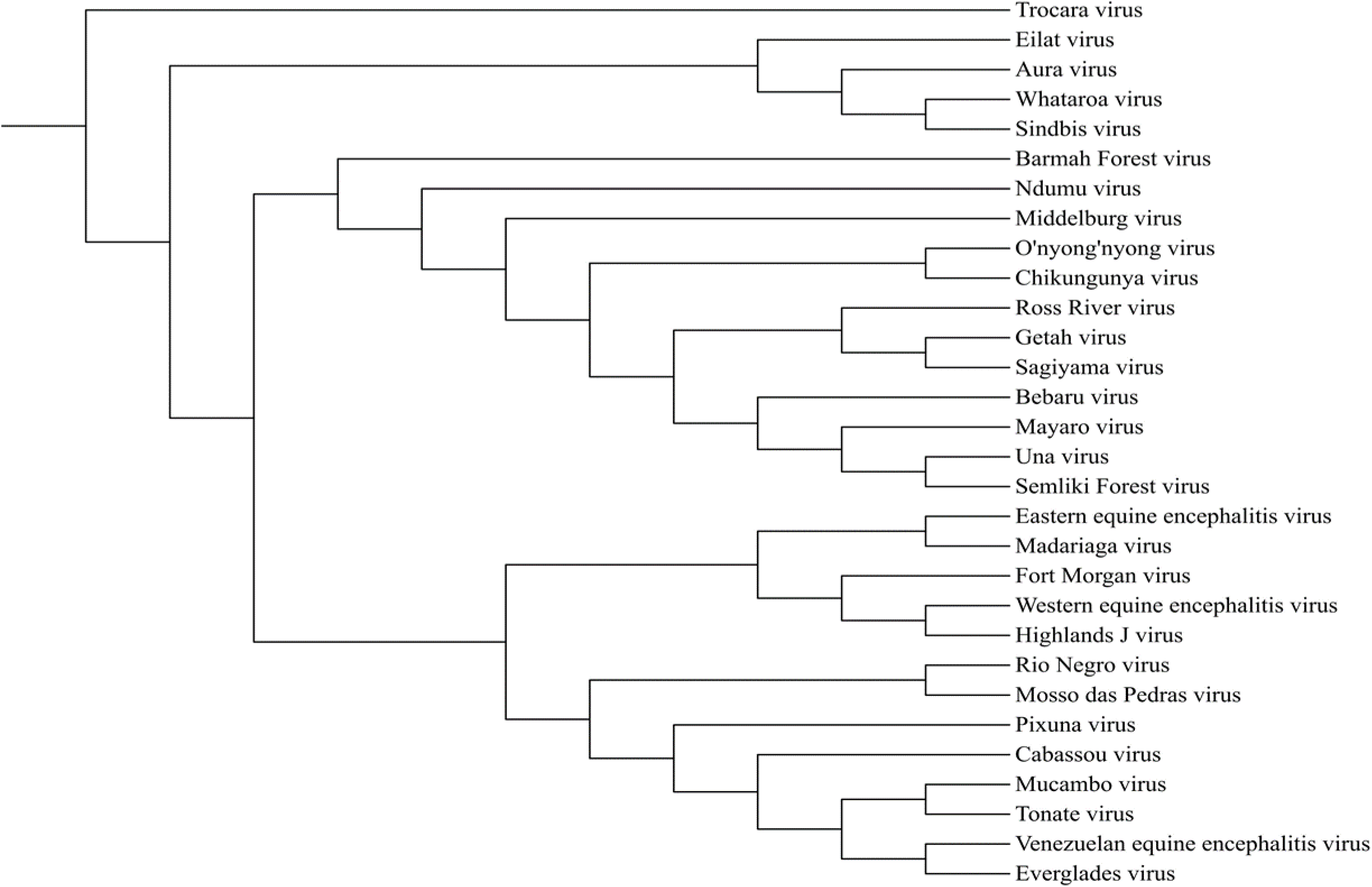

**Supplementary Figure 3.**
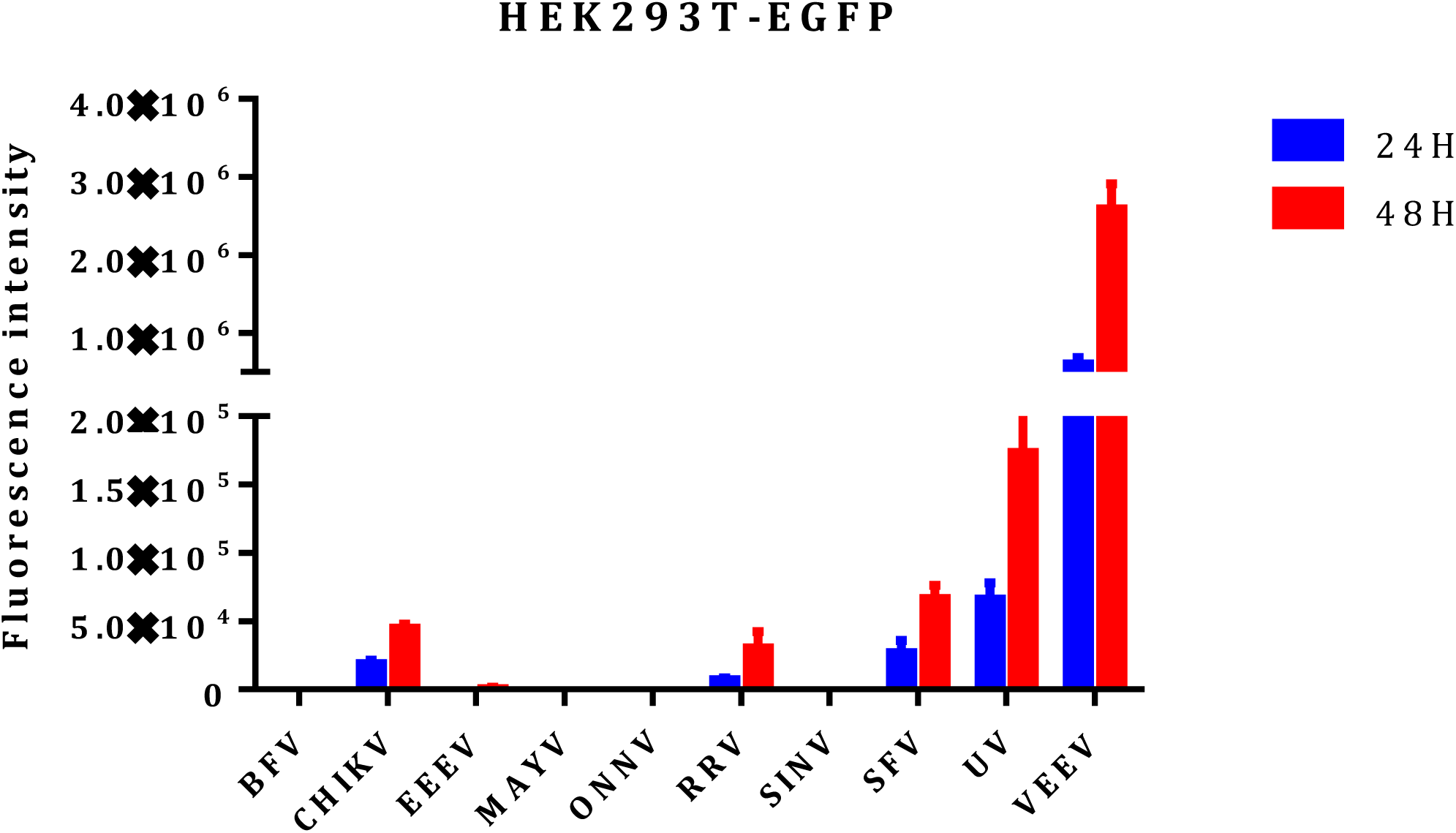

**Supplementary Figure 4.**
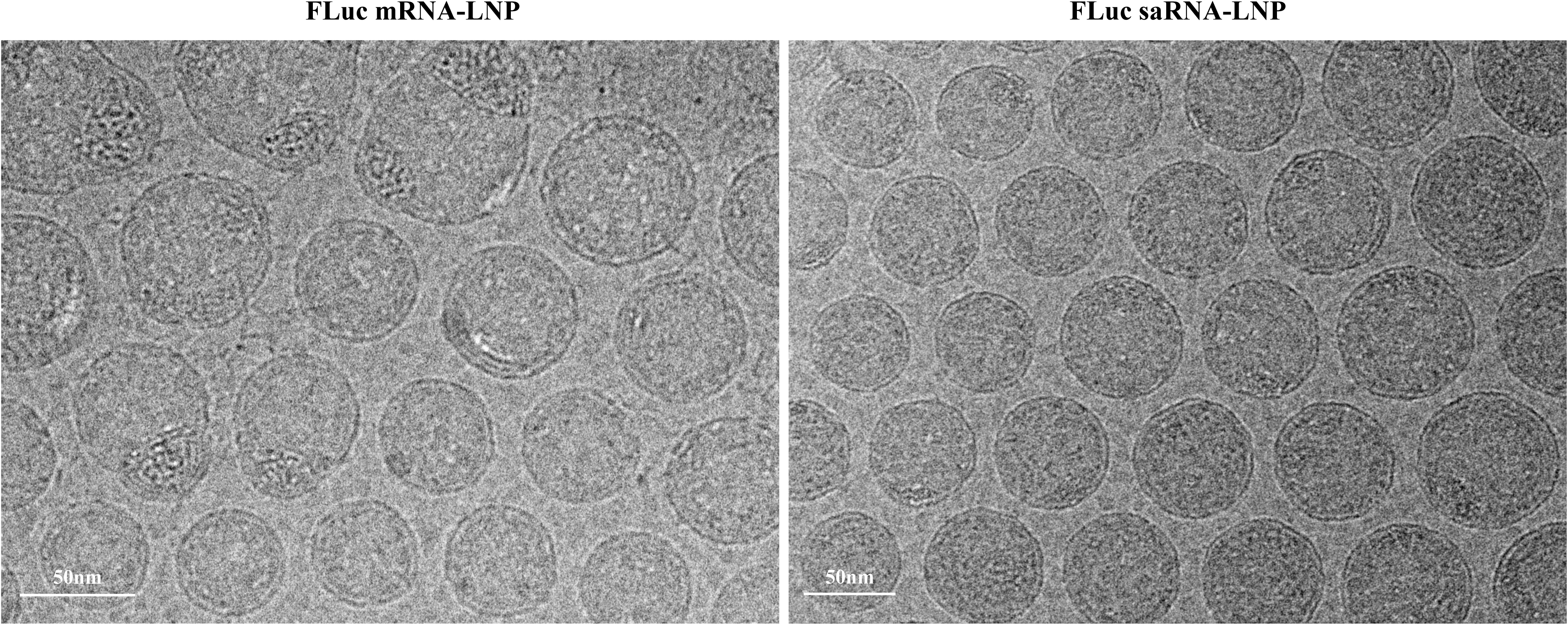

**Supplementary Figure 5.**
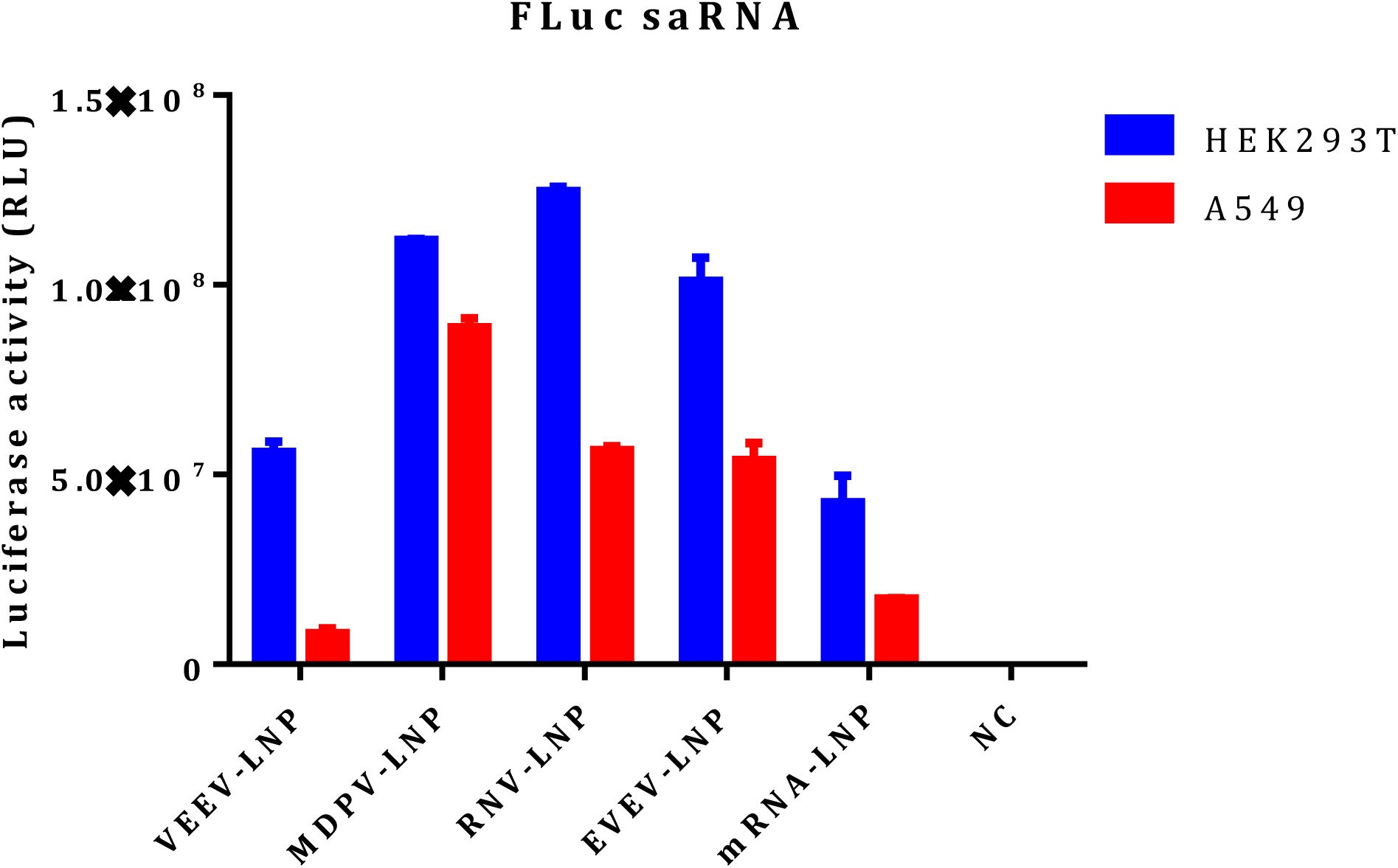

**Supplementary Table 1.**
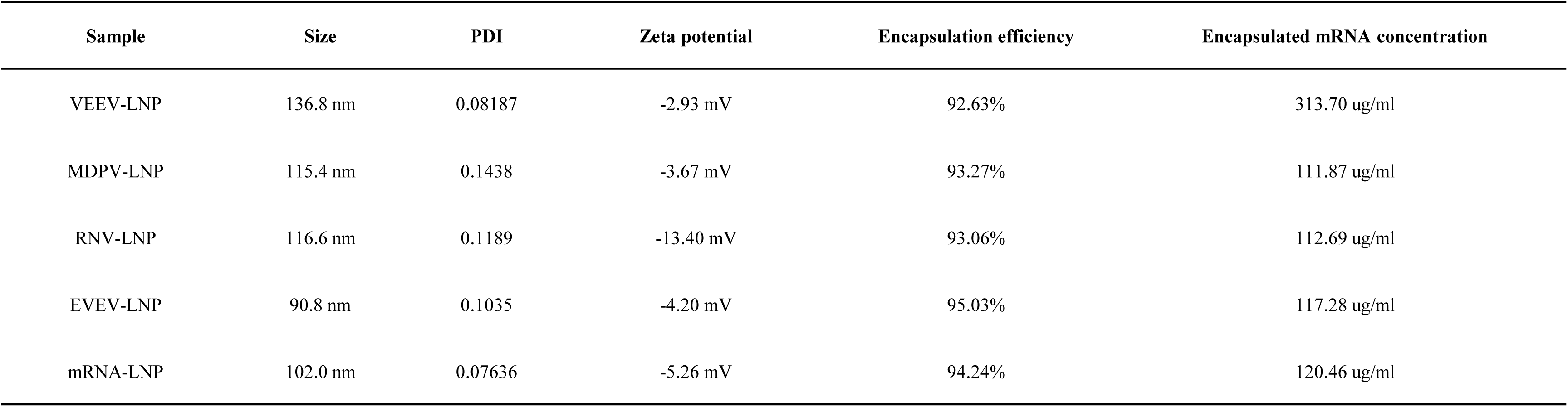

