## supplemental figure 1-5 for "Next-Generation saRNA Platforms: Systematic Screening and Engineering Enhances Superior Protein Expression and Organ-Specific Targeting for RNA Therapeutics"

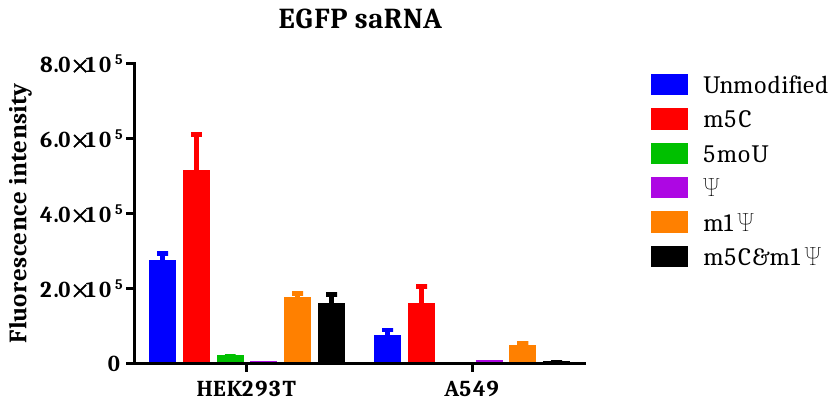

**Supplementary Figure 1.** Impact of nucleotide modifications on saRNA expression. HEK293T and A549 cells were transfected with 100ng different base modified EGFP saRNAs using lipofectamine. EGFP expression were determined 24h after transfection. Data shown as mean + SD of 3 independent experiments.

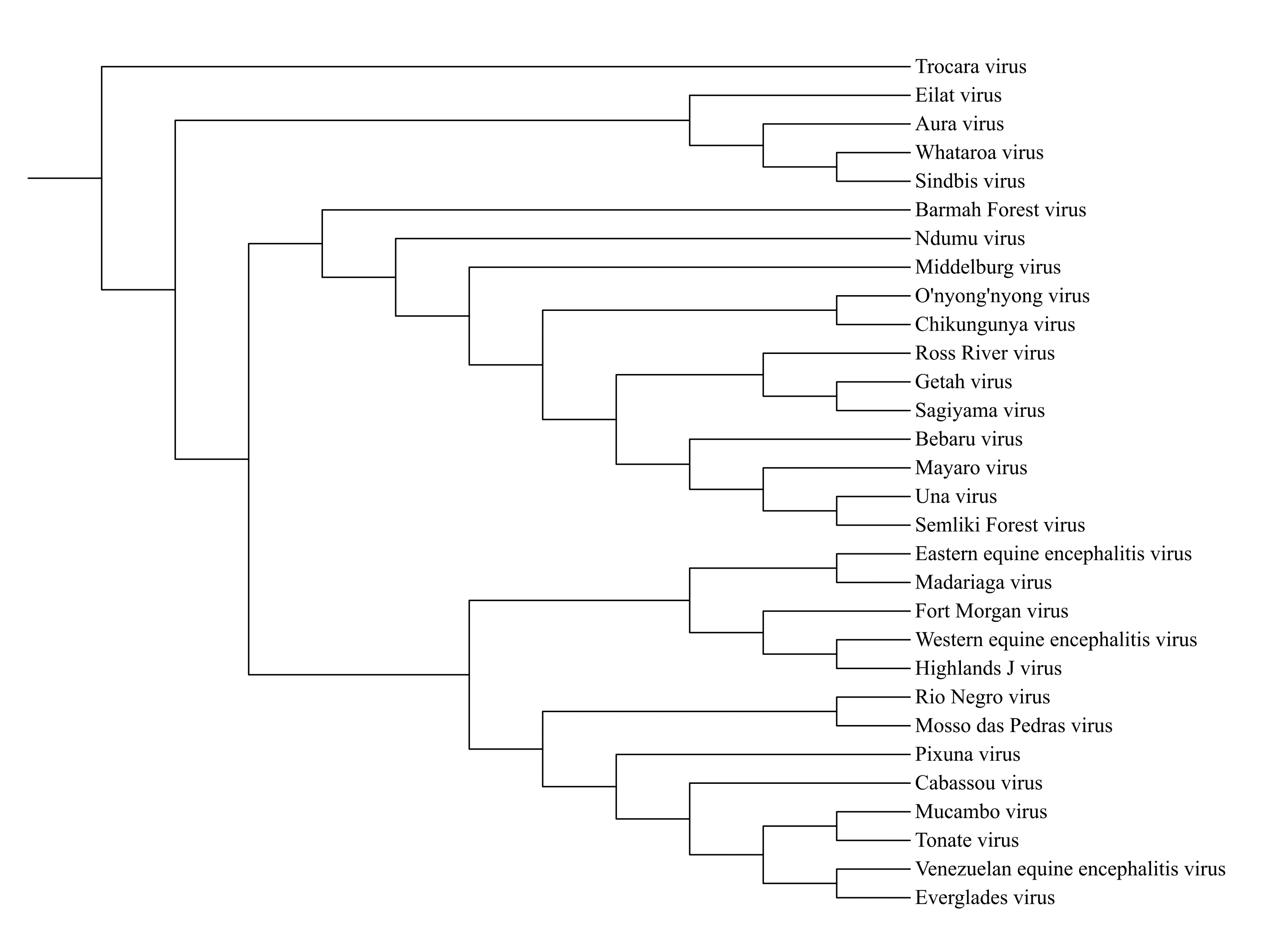

**Supplementary Figure 2.** Schematic representation of the phylogenetic tree of alphaviruses used for saRNA screening in this study.

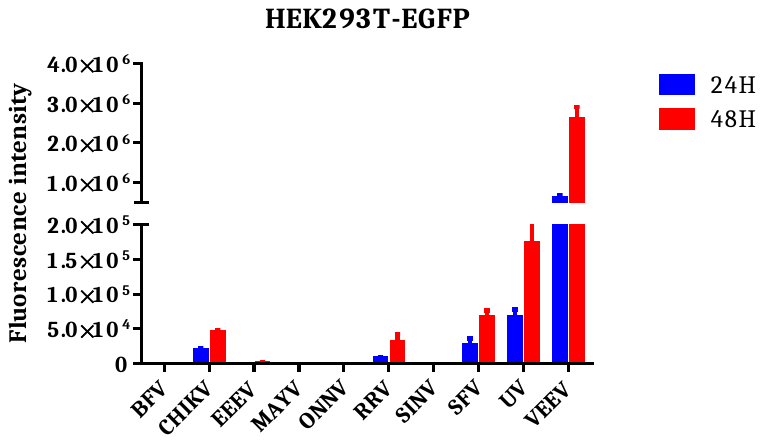

**Supplementary Figure 3.** Systematic in vitro screening of alphavirus-derived saRNAs. HEK293T cells were transfected with 10ng EGFP saRNAs using lipofectamine. EGFP expression were determined 24h and 48h after transfection. Data shown as mean + SD of 3 independent experiments.

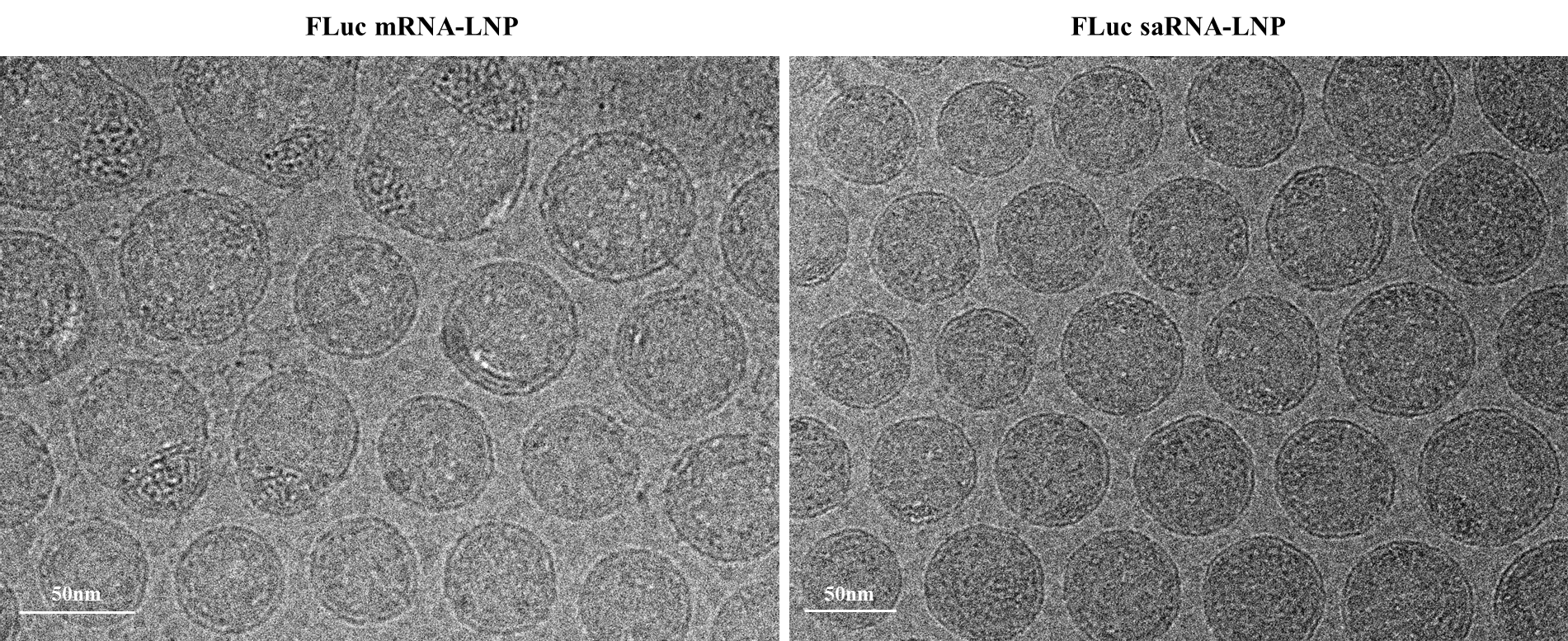

**Supplementary Figure 4.** Representative cryo-electron microscopy (cryo-EM) images of FLuc mRNA-LNP and FLuc saRNA-LNP formulations.

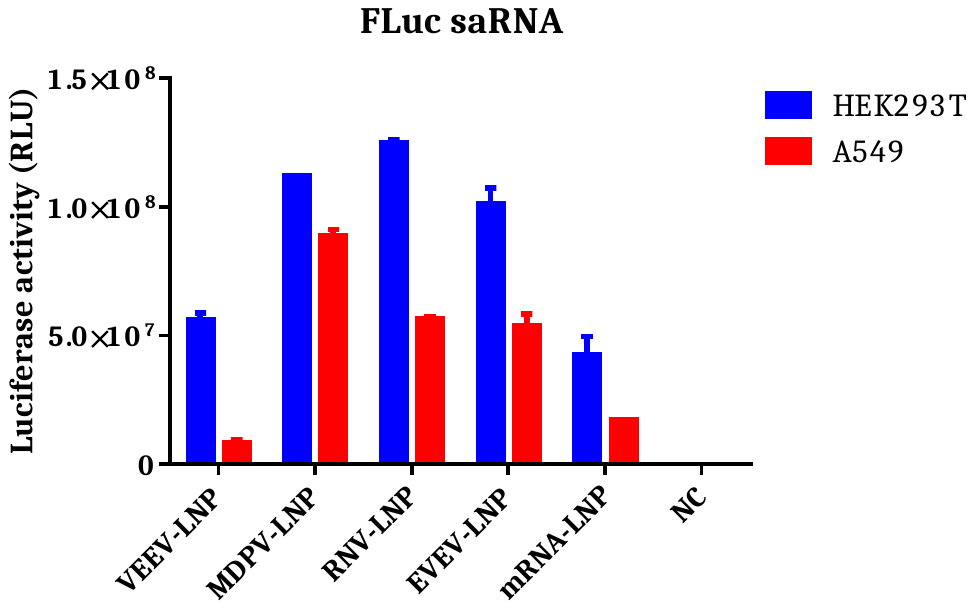

**Supplementary Figure 5.** Expression validation of FLuc saRNA-LNPs in HEK293T and A549 cells. HEK293T and A549 cells were transfected with 10ng FLuc saRNA-LNP or 100ng FLuc mRNA-LNP. Luciferase expression were determined 24h after transfection. Data shown as mean + SD of 3 independent experiments.

**Supplementary Table 1.** Physicochemical characterization results of the FLuc saRNA-LNP and mRNA-LNP formulations.

| **Sample** | **Size** | **PDI** | **Zeta potential** | **Encapsulation efficiency** | **Encapsulated mRNA concentration** |
| --- | --- | --- | --- | --- | --- |
| VEEV-LNP | 136.8 nm | 0.08187 | -2.93 mV | 92.63% | 313.70 ug/ml |
| MDPV-LNP | 115.4 nm | 0.1438 | -3.67 mV | 93.27% | 111.87 ug/ml |
| RNV-LNP | 116.6 nm | 0.1189 | -13.40 mV | 93.06% | 112.69 ug/ml |
| EVEV-LNP | 90.8 nm | 0.1035 | -4.20 mV | 95.03% | 117.28 ug/ml |
| mRNA-LNP | 102.0 nm | 0.07636 | -5.26 mV | 94.24% | 120.46 ug/ml |
